# Can we use anti-predator behavior theory to predict wildlife responses to high-speed vehicles?

**DOI:** 10.1101/2021.11.29.470451

**Authors:** Ryan B. Lunn, Bradley F. Blackwel, Travis L. DeVault, Esteban Fernández-Juricic

## Abstract

Animals seem to rely on antipredator behavior to avoid vehicle collisions. There is an extensive body of antipredator behavior theory that have been used to predict the distance/time animals should escape from predators. These models have also been used to guide empirical research on escape behavior from vehicles. However, little is known as to whether antipredator behavior models are appropriate to apply to an approaching high-speed vehicle. We addressed this gap by (a) providing an overview of the main hypothesis and predictions of different antipredator behavior models via a literature review, (b) exploring whether these models can generate *quantitative* predictions on escape distance when parameterized with empirical data from the literature, and (c) evaluating their sensitivity to vehicle approach speed via a simulation approach where we assessed model performance based on changes in effect size with variations in the slope of the flight initiation distance (FID) vs. approach speed relationship. We used literature on birds for goals (b) and (c). We considered the following eight models: the economic escape model, Blumstein’s economic escape model, the optimal escape model, the perceptual limit hypothesis, the visual cue model, the flush early and avoid the rush (FEAR) hypothesis, the looming stimulus hypothesis, and the Bayesian model of escape behavior. We were able to generate *quantitative* predictions about escape distances with the last five models. However, we were only able to assess sensitivity to vehicle approach speed for the last three models. The FEAR hypothesis is most sensitive to high-speed vehicles when the species follows the spatial (FID remains constant as speed increases) and the temporal margin of safety (FID increases with an increase in speed) rules of escape. The looming stimulus effect hypothesis reached small to intermediate levels of sensitivity to high-speed vehicles when a species follows the delayed margin of safety (FID decreases with an increase in speed). The Bayesian optimal escape model reached intermediate levels of sensitivity to approach speed across all escape rules (spatial, temporal, delayed margins of safety) but only for larger (> 1 kg) species, but was not sensitive to speed for smaller species. Overall, no single antipredator behavior model could characterize all different types of escape responses relative to vehicle approach speed but some models showed some levels of sensitivity for certain rules of escape. We derive some applied applications of our finding by suggesting the estimation of critical vehicle approach speeds for managing populations that are especially susceptible to road mortality. Overall, we recommend that new escape behavior models specifically tailored to high-speeds vehicles should be developed to better predict quantitatively the responses of animals to an increase in the frequency of cars, airplanes, drones, etc. they will be facing in the next decade.

## Introduction

Collisions between animals and vehicles (e.g., cars and airplanes) cause substantial economic damage, present a safety hazard to motorists, and are a source of mortality for wildlife [1–3]. In the United States, approximately 212,970 aircraft-bird collisions (2000-2020; hereafter, bird-strikes) cost the airline industry 594 million dollars, or 12,217 collisions annually representing at minimum US$34.65 million in cost per year [4]. In the United States, cars alone are estimated to kill 88.7 to 339.8 million birds annually [5, 6]. Reducing animal collisions with vehicles has both ecosystem and financial benefits.

One approach to address the animal-vehicle collision problem is to identify the behavioral mechanisms involved in animal decision-making when confronted with an approaching vehicle [6]. Formalizing those mechanism in models could eventually allow us to predict the responses of different species to vehicles and be used to manage the risk of collision [7]. However, to our knowledge there is no body of theory that has been explicitly developed to understand and predict interactions between animals and high-speed vehicles.

Antipredator behavior theory, which explains the responses of prey to predators, could potentially fill this theoretical gap [8]. However, to apply these models a key is that animals respond similarly to vehicle approaches as predator approaches. This has been empirically corroborated in the context of both car approaches and aircraft approaches (Blackwell et al.

2019b) [9–13]. Specifically, when animals are approached by vehicles, they become alert and later engage in escape behavior to avoid a collision [9, 14–16] .

There is an extensive body of antipredator behavior theory including several models of escape behavior that have been used extensively to predict the distance or time at which animals should escape from an approaching predator [8, 17]. However, vehicles are fundamentally different from predators in their approach speed, size, and degree of variation in approach trajectory [3] .

For instance, models of escape behavior generally predict that faster predator approach speeds result in an animal escaping at longer distances [18, 19]. This prediction has been empirically corroborated, primarily with human approaches during which experimenters either walk or jog towards a focal animal, which has obvious limitations in approach speed relative to common vehicle speeds [20–23]. Cars often travel in excess of 120 km/h on modern highways and large jet aircraft can travel up to 240 km/h during takeoff and landing. Yet all of the antipredator behavior models have been developed within the context of predator approach speeds. DeVault et al. (2015) [15] found that Brown-headed Cowbirds (*Molothrus ater*) were not able to avoid a simulated vehicle approach travelling faster than 120 km/h, suggesting that vehicle approach speed is a critical factor in determining whether animals are able avoid an impending collision.

Therefore, it is unclear whether current models of escape behavior, constructed with predator approach speeds, are appropriate to make predictions about when an animal should escape from a high-speed approaching vehicle.

Our overall goal was to assess whether existing models of antipredator behavior could be used in the context of animal-vehicle collisions. First, we reviewed the literature for different antipredator behavior models that make predictions about the distance (or timing) of escape decisions of a stationary animal upon the approach of a predator. Second, we determined whether each of these models could generate *quantitative* predictions about the distance at which animals escape, which is a common metric shared across models. To assess the ability of different models to generate quantitative estimates of escape distance, we parameterized them with published data [15]. Third, we explored whether those models that can generate quantitative predictions are sensitive to variations in the speed of a modern vehicle. Given the limited empirical data on behavioral responses to high-speed vehicles, we generated data with a simulation parameterized with empirical data from different bird species. These simulated data allowed us to establish whether models can predict changes in the distance at which animals escape with variations in vehicle speed relative to different behavioral rules animals use to escape when approached by threats (spatial margin of safety, temporal margin of safety, and delayed margin of safety).

Our focus was on quantitative rather than the qualitative predictions of these models because they hold a greater potential to improve the management of animal-vehicle interactions. Many of the antipredator behavior models yield qualitative predictions in terms of flight initiation distance (i.e., the distance between the threat and the animal when the latter escapes; hereafter FID) and there are extensive empirical estimates of FID from multiple vertebrate taxa [19]. Consequently, antipredator behavior models that can generate species-specific quantitative FID predictions relative to high-speed vehicle approaches could aid conservation practitioners and stakeholders in identifying vulnerable species and areas (existing and future roadways, airports, etc.), estimate the mortality risk for both wildlife and humans, explore the ecological consequences of potential management practices aimed at mitigating the rate of animal-vehicle collisions, and forecast the economic consequences of animal-vehicle collisions (e.g., insurance claim estimates, property damage).

## Methods

### General overview

Our study is divided in three main sections that we explore for each model: (1) overview of the main hypothesis and predictions, (2) ability to generate quantitative predictions on escape distance based on the speed of the approaching threat, and (3) sensitivity to variations in vehicle approach speed. In (1), we reviewed the literature to identify different models of escape behavior that were potentially applicable to a scenario where an immobile prey is approached by a high speed vehicle. In (2), we attempted to generate quantitative predictions for each model with empirical data from DeVault et al. 2015 [15] which measured the widest range of approach speeds under standardized conditions. In (3), we examined only models from which we were able to generate quantitative predictions by simulating data based on parameters that we extracted from another literature review. With the simulated data, we assessed the variation in the ability of each model to predict escape distance with different approach speeds by estimating the effect sizes. In the following sections, we describe in detail our methodology for each step.

### Identifying anti-predator escape behavior models

We only reviewed models of antipredator escape behavior that represented the scenario of a stationary animal being approached by predator, because it represents a common scenario in animal-vehicle interactions. An observational study found that 96% of birds sampled at an airport were stationary at the beginning of the interaction with an aircraft [15]. Of the stationary birds, 62% were on the ground and 38% were perching on structures at different heights [15]. Our initial review of the literature began by identifying anti-predator escape behavior models reviewed in Cooper & Blumstein’s (2015) book [8]. Of the nine models of predator escape behavior reviewed in chapter 2, we identified three which potentially could be applied to animal-vehicle interactions: economic escape model [18], Blumstein’s economic escape model [24], and optimal escape model [25]. Next, we used both Google Scholar and Web of Science to search for additional papers. We searched for papers from 1986 (the year of publication of Ydenberg and Dill’s seminal paper) to 2020. We focused our search on the following keywords: FID, flight initiation distance, approach speed, model, escape, and antipredator escape behavior.

We identified the following eight models: the economic escape model [18], Blumstein’s economic escape model [24], the optimal escape model [25], the perceptual limit hypothesis [26], the flush early and avoid the rush (FEAR) hypothesis [27], the looming stimulus hypothesis [28–29], the visual cue model [30], and a Bayesian model of escape behavior [31]. For each model, we limited our overview in the text to the main hypotheses and predictions for the sake of space. However, a detailed account of all the hypotheses, predictions, and assumptions are provided in S1. Our literature search yielded one more model (Broom and Ruxton’s (2005) escape behavior for cryptic prey) [32] that we decided not to include because in the model the distance at which an animal escapes is dependent on changes in the behavior of the approaching predator, which is not easily translated into the context of an approaching vehicle.

The eight models used specific terms to define the different stages of a predator-prey encounter. We closely followed terminology from Cooper and Blumstein (2015) [8] and adjusted the terms to a scenario with vehicles to standardize the discussion between models. Starting distance (SD) was considered the distance between the animal and the vehicle when the latter begins the approach. Detection distance (DD) was considered the distance between the animal and the vehicle when the former detects the vehicle. Alert distance (AD) was considered the distance between the animal and the vehicle when the former begins showing alert behaviors (e.g., head-up posture). Flight initiation distance (FID) was considered the distance between the animal and the vehicle when the former initiates escape. Time to collision (TTC) is how much time away the approaching vehicle is from the animal (i.e., a collision occurs when TTC = 0 if the animal does not escape in a timely fashion).

### Can models generate quantitative predictions?

A major component of using antipredator escape behavior models to understand animal-vehicle interactions is establishing their ability to make quantitative predictions. Our first step for each model was to use empirical data from the literature to parameterize each model. To that end, we needed published data on the responses of animals to vehicles (rather than predators or humans) travelling at known speeds. Unfortunately, little empirical data were available to feed the different parameters each model requires relative to responses to vehicles [9, 14–16]. We chose to parameterize the different models with data from DeVault et al. [15] because the study features the largest range of approach speeds to which an animal was exposed in a systematic and experimental manner. In that study, brown-headed cowbirds were exposed to virtual approaches of a vehicle at different speeds (60, 90, 120, 150, 180, 210, 240, and 360 km/h) [15].

Whenever possible, we attempted to infer or directly use parameters from the data obtained from DeVault et al. (2015) [15] in every model that allowed us to do so. The purpose of attempting to make predictions from a common dataset was to maximize our ability to make the predictions comparable across models. Besides approach speed, some models required some extra parameters, which were derived from the literature on brown-headed cowbirds or phylogenetically related species [33–38]. We explain the source of these parameters in the section for the corresponding model to facilitate the assessment of our rationale. We made the following assumptions throughout: (a) the vehicle approaches an animal at a constant speed and (b) the vehicle approaches an animal directly (i.e., the approach angle is equal to 0) and moves in a straight, undeviating path. All quantitative predictions were generated using R 4.0.2 (R Core Team, 2020) [39], and we made the data and code available for this section in S5.

### Are models sensitive to vehicle approach speed?

We assessed the degree to which the models that can generate quantitative predictions are sensitive to vehicle approach speed. Five of the eight models reviewed were capable of making quantitative predictions (see Results), but we only assessed three of them relative to sensitivity to speed: the FEAR hypothesis, the looming stimulus hypothesis, and the Bayesian optimal escape model. Two of the five models (perceptual limits hypothesis, visual cue model) do not incorporate the speed of the approaching threat in their mathematical formulation; therefore, we could not evaluate them. The looming stimulus hypothesis and the Bayesian optimal escape model both explicitly incorporate a parameter establishing the approach speed of the threat [29, 31]. Although the FEAR hypothesis does not explicitly incorporate a parameter to account for approach speed [27, 40] ; we were able to evaluate the effects of speed indirectly by considering how phi index values, a proxy for effect size (i.e., the magnitude of an effect) [41], changes depending on how an animal adjusts FID with approach speed [42].

The core of our approach considered the relationship between FID and the speed of the approaching threat. However, different species have different FID responses to approach speed [19,20,40,43,44]. Consequently, we focused on the *slope* of the FID vs. approach speed relationship (Fig 1), which allowed us to capture the degree of interspecific variation in response to approach speed. Positive slopes indicate an increase in FID as approach speed increases, slopes close to zero indicate that FID does not change as approach speed increases, and negative slopes indicates that FID decreases as approach speed increases. Considering the slope of the FID vs. approach speed relationship also gives the opportunity to associate these patterns with different escape rules animals exhibit when escaping from approaching threats: the temporal margin of safety [45, 46], the spatial margin of safety [45–48], and another phenomenon that we call here the delayed margin of safety (see below).

**Figure 1.**
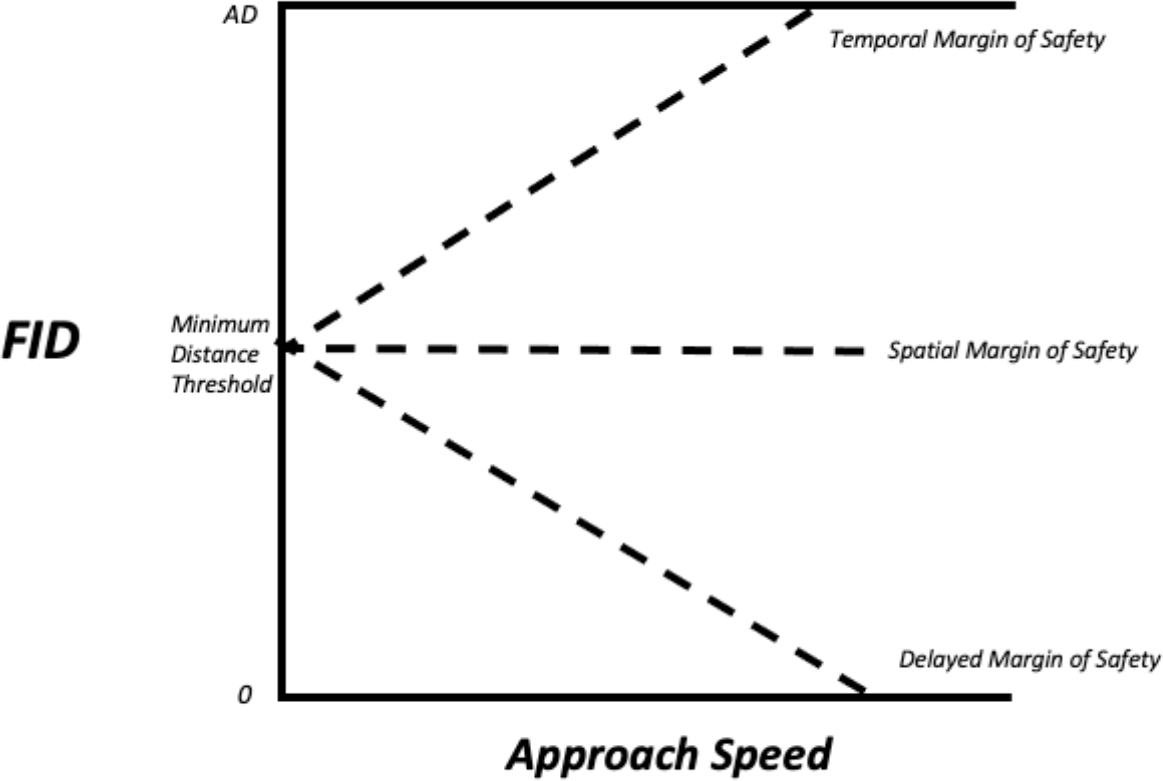
The different escape rules different species may follow based on the slope of the relationship between FID and approach speed. Positive slopes are associated with the temporal margin of safety, a slope near zero is associated with a spatial margin of safety, and negative slopes are associated with the delayed margin of safety. If the slope is positive FID increases until it reaches AD, at which point FID is equal to AD. If the slope is negative FID decreases until it reaches 0, at which point FID is equal to 0. The minimum distance threshold is the last distance at which the animal will tolerate the approach of a threat whereafter if the threat continues to approach to where the animal will escape, regardless of the threats approach speed.

Animals use the temporal margin of safety when individuals associate risk with *time* away from a potential collision with an approaching threat. Individuals following the temporal margin of safety maintain a set amount of time away from the approaching threat for escape [15,45,46] . Consequently, as approach speed increases, FID is expected to increase to maintain the same amount of time away from the threat, leading to a positive slope in the relationship between FID and approach speed.

Animals use the spatial margin of safety when individuals associate risk with *distance* away from a threat. Individuals following the spatial margin of safety maintain a set distance away from the approaching threat for escape, regardless of the speed of the oncoming threat [15,45,46]. Threats approaching at different speeds cover the exact same distance to reach the same location but at different times. Consequently, as approach speed increases, FID is expected not to vary with speed, leading to a slope close to 0 in the relationship between FID and approach speed.

The delayed margin of safety intends to reflect a phenomenon observed in some papers in which the relationship between FID and speed is negative (as opposed to the predictions of the temporal and spatial margins of safety). The delayed margin of safety could be the result of individuals delaying escape upon detecting an approaching threat. Species following the delayed margin of safety might be compromised to detect and track a threat approaching because their limited attention is allocated to monitoring some other pertinent stimuli (i.e., foraging, conspecifics, etc.) and the potential approaching threat [49–53]. This division of attention can result in the continued approach by the threat, thereby decreasing the distance between the animal and the threat without the former escaping (i.e., the distracted prey hypothesis) [27, 54]. The pattern of shortening the distance between the animal and the vehicle can be more pronounced as the speed of the vehicle increases. Consequently, if escape is delayed, approaches at higher vehicle speeds would lead to a decrease in FID, resulting in a negative slope in the relationship between FID and approach speed.

We used five steps to evaluate sensitivity to speed (Fig 2). Ultimately (step 5), we established the degree of sensitivity to approach speed for each of the three models (FEAR hypothesis, looming stimulus hypothesis, Bayesian optimal escape) by assessing the relationship between effect size and the slope of the FID vs. approach speed relationship (Fig 2). We estimated effect size as the amount of variation in FID accounted for by approach speed (see details for each model below). Because we explored effect size using different slopes, we could make inferences about the sensitivity to speed for different rules of escape (temporal margin of safety, spatial margin of safety, delayed margin of safety). We used a simulation approach for this portion of the study by generating datasets, based on parameters obtained from the literature, exploring the effects of the variation in approach speeds and slopes on FID (Fig 2). Simulation studies have been used extensively in ecology and evolution,[55–56] particularly when empirical data are lacking, like in our case. We focused on birds, as they have a high representation in the literature of animal-vehicle collisions,[9,12,15,16] which facilitated obtaining empirical estimates for the different parameters.

**Figure 2.**
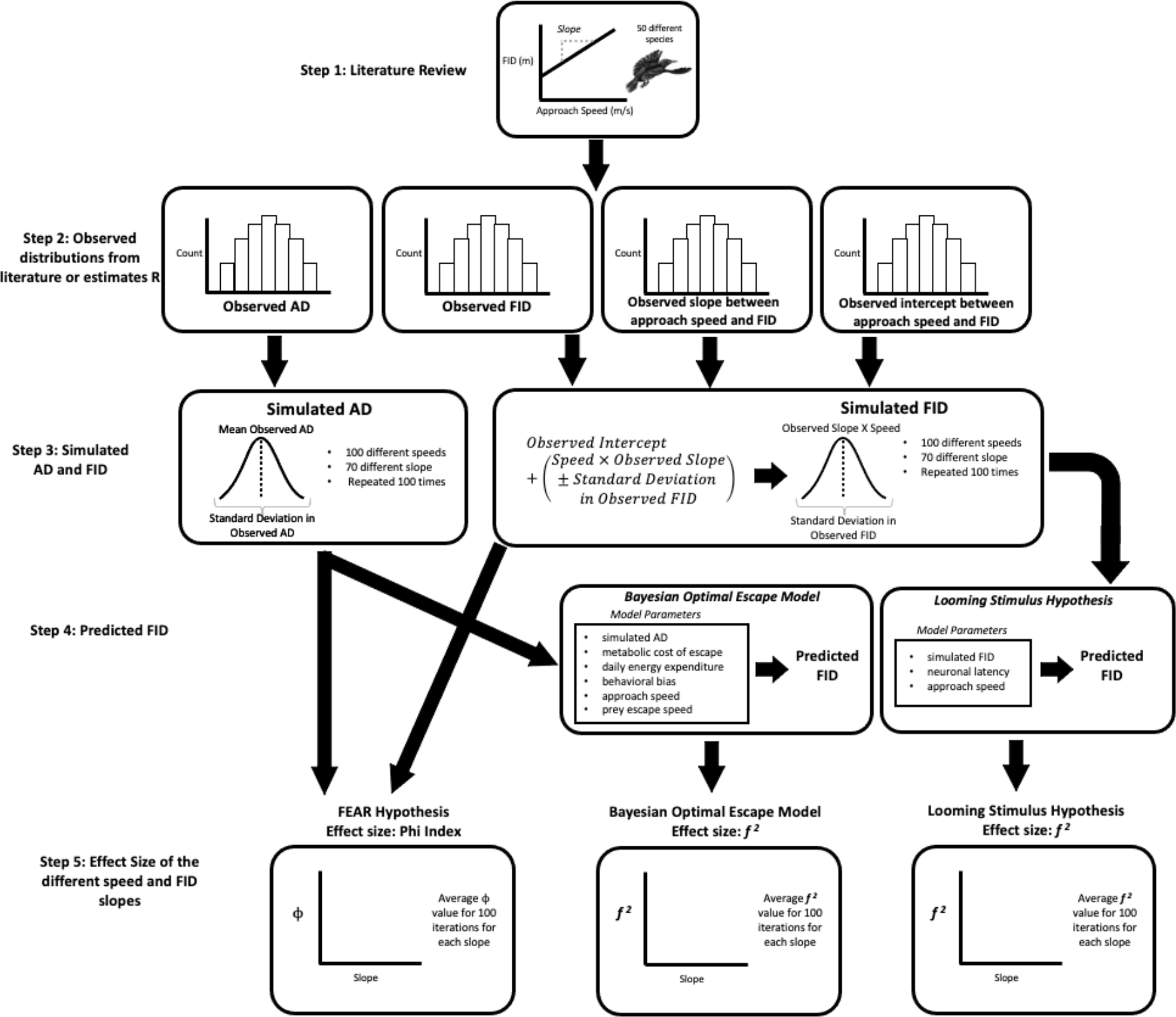
A flowchart with the different steps used to evaluate sensitivity to speed for the FEAR hypothesis, Looming stimulus hypothesis, and the Bayesian optimal escape model. Step 1: we reviewed the literature for studies which explored the effect of approach speed on FID in different avian taxa. The goal was to characterize the range of observed slopes of the FID vs. approach speed relationship when birds were exposed to human-related threats. Step 2: We established the distributions of AD, FID, then slope and intercept of the FID vs. approach speed relationship. Step 3: We simulated AD and FID based on the observed distributions from step 2 All three models required information on either FID, AD, or both to make quantitative predictions. Step 4: We generated model-specific FID predictions using additional parameters when needed. Step 5: We evaluated sensitivity to approach speed by estimating the average effect size of approach speed on model predicted FID at different slope values for each model.

In the first step, we again reviewed the literature for studies which explored the effect of approach speed on FID in different avian taxa. The goal was to characterize the range of observed slopes of the FID-vs.-approach-speed relationship when birds were exposed to human-related threats. In this case, we searched the literature using Google Scholar for studies published between 1950 through 2021 and that reported species-specific FID data at different approach speeds. We used the following keywords: speed, flight initiation distance, and birds.

Of the 7,190 studies returned, 9 met the criteria of explicitly assessing the effect of different approach speeds on FID in birds [14-16,57-62] (S2). From these nine papers, we found data for 50 different bird species that were exposed to a variety of different direct approach types (i.e., humans, bicycles, cars, and buses). We considered not only vehicle related approaches but also approaches by humans walking towards birds, to have the widest representation of species given the relatively low number of species (24) with published data on FID responses (at different vehicle speeds) to vehicle approaches. We only considered species that were directly approached.

We estimated the slope between FID (m) and approach speed (m/s) for each species with a linear regression. Most of the studies considered approach speed as a categorical factor with different numbers of levels. We took the mean FID value per speed treatment in those cases to accommodate our linear regressions. Some studies considered only two levels of approach speed, which could severely affect the estimate of the slope and intercept. Nevertheless, we decided to include them given the limited number of studies available and the fact that this exercise was oriented to defining a range of slopes rather than obtaining accurate species-specific slope estimates. All quantitative predictions and linear regressions were generated using R 4.0.2 [39]; these data and code are available for this section in S5.

In the second step, we established the distributions of AD, FID, slope and intercept of the FID vs. approach speed relationship. Each distribution consisted of 50 data points related to each species (Fig 2, S2). The AD and FID distributions consisted of the mean observed AD and FID for each species. Of the 9 studies we considered, AD was reported for only two of the 50 species. When AD data were not reported in the studies we identified (48 species), we used AD data reported for the same species from other papers (2 species) [63, 64] . If no AD measurements existed for a species, AD was estimated based on the species mean body mass (46 remaining species) (S2) following Blumstein et al. (2005) [65]. AD values ranged from 9.9 m to 63.7 m, with the mean (± SD) AD being 49.55 ± 10.51 m (S2). FID values ranged from 5.55 m to 128.38 m, with mean (± SD) FID being 43.97 ± 31.08 m (S2). The slope of the relationship between FID vs. speed ranged from -37.46 to 32.33 (mean ± SD, -2.12 ± 10.95; S2). The intercept of the relationship between FID vs. speed ranged from -75.70 to 128.4 (mean ± SD, 42.98 ± 46.27; S2). The four different distributions histograms are available in Fig S7. We considered all these parameters in developing our simulation.

In the third step, we generated AD and FID based on the observed distributions from step 2 (Fig 2). All three models required information on either FID or AD, or both to make quantitative predictions. The FEAR hypothesis considers the relationship between FID and AD, the looming stimulus hypothesis considers how a given FID is affected by neuronal latency, and the Bayesian optimal escape model considers AD [27,29,31,42]. Consequently, we generated pairs of AD and FID for each combination of approach speed (100 speeds) and slope (70 slopes). We selected a systematic change in approach speed from 1 to 100 m/s in 1 m/s intervals. The lower bound of approach speed was chosen because it corresponded to a typical human approach speed. The upper bound of approach speed was chosen because it corresponded with the speed of airplanes that some birds are exposed to [15]. We varied our slopes varied from -37 to 32 at intervals of 1 unit, which was based on step 2 (S2). We generated a pair of AD and FID at each approach speed and slope combination, which led to a total of 7,000 different pairs of values. We repeated the generation or simulation process 100 times to ensure that our results were not the result of a single simulation, which generated a total of 700,000 pairs of AD and FID values (step 3, Fig 2). AD and FID values were simulated with R. 4.0.2 (R Core Team, 2020 [39], S5).

Specifically when generating simulated AD data, we assumed that AD would not change with approach speed and slope because the studied models assumed that detection distance is a constant [18,24–31]. We simulated AD values from a normal distribution. The distribution was centered on a mean of 48.55 m with SD = 10.51 m based on step 2 (Fig 2). The simulated FID values changed with approach speed and slope. Our simulated FID data were based on the equation of the line, whereby FID = (slope x approach speed) *+* intercept. We kept the intercept constant because the intercept of the linear relationship between FID and approach speed (i.e., the FID when approach speed is 0) is by definition invariant with speed. The intercept can be interpreted as a minimum distance threshold at which if a threat approaches any closer the animal will escape regardless of the circumstances [24]. The implication is that species with negative slopes would still maintain a minimum distance away from a threat [24]. We simulated FID values from a normal distribution where the mean of our distribution was the product of approach speed and slope and the standard deviation was based on the mean standard deviation in FID among the 50 different species. Slope and approach speed were systematically varied.

Two steps were followed to create the equation used to simulate FID. First, we simulated FID values by multiplying a given slope (from -37 to 32) by a given approach speed (from 1 m/s to 100 m/s) that varied ± 31.08 m based on the observed SD in mean FID. Second, we added the mean intercept from the observed distribution of 50 species (42.98 m; step 3 Fig 2).

For our simulated FID values, we established boundaries, given the range of approach speeds and slopes we explored. An animal cannot escape from a threat if the threat has not been detected. FID is constrained by detection, which several models assume to be AD [27,30,31]; therefore, FID was less or equal to AD. The minimum value of FID was set to zero because FID cannot be negative [8]. The most an animal can delay escape is up until the point where the threat comes in contact with the animal (i.e., FID = 0). The implication is that when simulating negative slopes, as speed increases, FID continues to decrease only until it reaches a distance of 0 m.

In the fourth step, we generated model-specific FID predictions using additional parameters, when needed (step 4; Fig 2). The FEAR hypothesis only considers how FID changes relative to AD; therefore, it cannot generate predicted FID [27,40,42]. As a result, we skipped step four in the case of the FEAR hypothesis (Fig 2). The looming stimulus hypothesis explicitly considers neuronal latency along with approach speed and FID. We reviewed the literature for empirical data on neuronal latency [29, 66] for birds, but only found data for the Rock Pigeon (*Columba livia*). Neuronal latency in the pigeon ranged from 0.05 to 0.1 s [66]. For each combination of approach speed and slope we generated quantitative FID predictions with 25 different neuronal latency values spaced out in even intervals following the range reported in Wang & Frost (1992) [66]. As a result, we obtained 17,500,000 FID predicted values for the looming stimulus hypothesis (i.e., 25 neuronal latencies *\** 700,000 simulated FID values) (Fig 2).

The Bayesian optimal escape model considers several parameters when predicting FID: maximum escape velocity, approach speed, behavioral bias, AD, daily energy expenditure, and the metabolic cost of escape [66]. We did not have sufficient empirical information to estimate the parameter for behavioral bias, *α*; therefore, we assumed no bias (*α* = 0.5) [15, 31]. Maximum escape velocity, the metabolic cost of escape, and daily energy expenditure can all be estimated as a function of a species body mass [34,35,37]. Body mass values were based on the species mean (male and female) of the 50 different bird species studied [36]: which ranged from 11.5 grams (Superb Fairy Wren, *Malurus cyaneus*) to 6.22 kilograms (Black Swan, *Cygnus atratus*). The distribution of body masses was positively skewed; consequently, we first log transformed body mass values and then sampled 50 different values (see below) in even intervals. We chose 50 different values based on the 50 different species considered in the review. The inverse log of those values produced a distribution which represented the empirically observed distribution of body masses. As a result, we generated 35,000,000 FID predicted values for the Bayesian optimal escape model (i.e., 50 body mass values *\** 700,000 simulated FID values) (step 4; Fig 2).

In the fifth step, we evaluated sensitivity to approach speed by estimating the average effect size of approach speed on model predicted FID at different slope values for each model. We evaluated the FEAR hypothesis with a proxy of effect size: the phi index or *Φ* (step 5, Fig 2). *Φ* assesses how well the AD and FID relationship fits a line with a slope of 1 when the intercept is zero, ranging from 0 to 1. When *Φ* = 0, both AD and FID are equal to 0 because when AD, the predictor of FID according to the FEAR hypothesis, is equal to 0, the assumption is that AD ≥ FID, which limits FID to 0. *Φ* > 0 but *<* 0.5 suggest that animals do not follow the FEAR hypothesis. *Φ* ≥ 0.5 indicate that animals follow the FEAR hypothesis. We evaluated the sensitivity to approach speed of the looming hypothesis and the Bayesian optimal escape model by using the *f ^2^* effect size, which was calculated as: 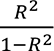[41] (step 5, Fig 2). We estimated *R^2^* from linear regressions between the predicted FID vs. approach speed. *f ^2^* values were categorized as small (0.02), medium (0.15), or large (0.35) based on Cohen (1992) [41] recommendation for *f ^2^* values. We estimated effect size values for each slope value by taking the mean phi index (FEAR hypothesis) or *f ^2^* (looming stimulus hypothesis and Bayesian optimal escape model) across all approach speeds and slopes.

We presented our results following two different approaches due to the different number of parameters each model considers (step 5, Fig 2). The FEAR hypothesis only considers AD and FID parameters, so a 2-dimensional representation of the average effect size relative to the 70 different slopes of the FID vs. approach speed was sufficient. However, both the looming stimulus hypothesis and the Bayesian optimal escape model consider additional parameters; therefore, we generated for these models 2-dimensional representations similar to the ones for the FEAR hypothesis as well as 3-dimensional representations considering the extra parameters (step 5, Fig 2). For the looming stimulus hypothesis, the 2-dimensional representation between effect size and slope used the mean neuronal latency value (0.075) [66] but we included neuronal latency as a third axis in the 3-dimensional representation (in even intervals from 0.05 s to 0.1 s). The Bayesian optimal escape model also considers maximum escape velocity, daily energy expenditure, and metabolic cost of escape, which can be estimated based on body mass. Thus, we summarized these parameters by using body mass as a proxy in our representation of this model. For the Bayesian optimal escape model, the 2-dimensional representation between effect size and slope used the mean body mass of our 50 studied species (267.4 g), but we included body mass as a third axis in the 3-dimensional representation (in even intervals on a log scale from 11.5 g to 6.22 kg).

We interpreted the results in terms of how effect size changes for three slope regions related to the different escape rules. The three slope regions are positive slopes from 1 to 32 (temporal margin of safety), slope = 0 (spatial margin of safety), and negative slopes from -1 to - 37 (delayed margin of safety). For the spatial margin of safety, FID is expected to be invariant with approach speed [15,46,48]. This means that the slope of the FID vs. approach speed relationship would not be significantly different from 0 in the context of a linear regression.

However, operationally, the challenge was to define the range of values around 0 to be considered different from 0 under the spatial margin of safety. To address that issue, we generated yet another set of simulated AD and FID at different approach speeds and slopes.

Similar to the previously described simulation, simulated AD data did not change with approach speed and simulated FID data were based on the equation of a line whereby FID = (slope *\** approach speed) *+* intercept. We kept the intercept constant because the intercept of the linear relationship between FID and approach speed (i.e., the FID when approach speed is 0) is by definition invariant with speed. The lower bound of the simulated FID values was 0, and the upper bound was defined by AD. Four steps were followed to generate AD and FID values.

First, we generated AD data from a normal distribution centered on a mean AD of 50 m with a standard deviation of 10 m. Second, we generated FID data from a normal distribution, where the mean was determined by multiplying slope and approach speed. We used 8 different approach speed treatments (60 km/h, 90 km/h, 120 km/h, 150 km/h, 180 km/h, 210 km/h, 240 km/h, 360 km/h). These approach speeds composed the largest range of different approach speeds where an animal was reported to follow a spatial margin of safety from 60 km/h to 150 km/h; however, no specific behavioral rule was observed at speeds faster than 180 km/h [15].

Without any prior expectation of which slope values might result in a non-significant model we used two different sets of simulated slopes. The first set ranged from – 1 to 1 in even increments of 0.05 and then the second set ranged from -10 to 10 in even increments of 0.5. FID values had a standard deviation of 10 m identical to the variation in AD. Third, we established that the intercept of the line was 25 m because it was half the distance of the mean alert distance. Fourth, we generated FID by multiplying slope values (i.e., -1 to 1, by 0.05 or -10 to 10 by 0.5) with approach speed (i.e., 60 km/h, 90 km/h, 120 km/h, 150 km/h, 180 km/h, 210 km/h, 240 km/h, 360 km/h) from step 2 and then adding the intercept of 25 m from step 3.

Each run of the simulation generated 20 FID data points for the 8 different approach speed treatments (160 total FID data points) similar to DeVault et al. (2015) [15] for each slope, which led to a total of 6,560 different FID values. We repeated both simulations 500 times, which generated a total of 3,280,000 FID values. To evaluate which slope values adhered to the spatial margin of safety we estimated the mean *p-value* for each slope across 500 simulation iterations with a linear regression between FID and approach speed. Approach speed was treated as a categorical variable because seldomly in controlled environmental studies can different approach speeds be treated as a continuous variable. Our simulation suggests that based on the empirical data from DeVault et al. (2015) [15] that a spatial margin of safety had a slope between - 0.1 and 0.1. (S3).

## Results

### Economic escape model

#### 1. Overview

The Economic escape model proposes an ultimate explanation because it assumes that the distance at which an animal escapes will affect fitness. Ydenberg and Dill’s (1986) [18] graphical model represents a scenario in which an animal monitors an approaching predator and escapes at the distance which minimizes the cost to future fitness (S1.1). In the model, the interaction between predator and prey begins at the point where the predator starts its approach (SD).

Therefore, the model assumes that at some distance (unspecified by the model, possibly >SD) the predator is detected and that the approach is monitored by the prey. The model posits that the decision of the animal to escape at a certain distance is a function of two cost curves: the cost of fleeing and the cost of not fleeing (Fig 3a). The cost of fleeing curve is associated with the opportunities an animal foregoes when fleeing from a predator (i.e., opportunity cost), and this curve increases with distance from the predator. If the animal escapes prematurely, it will forego a larger amount of benefits (i.e., foraging) than if it delays escape (Fig 3a – blue solid line) [18].

**Figure 3.**
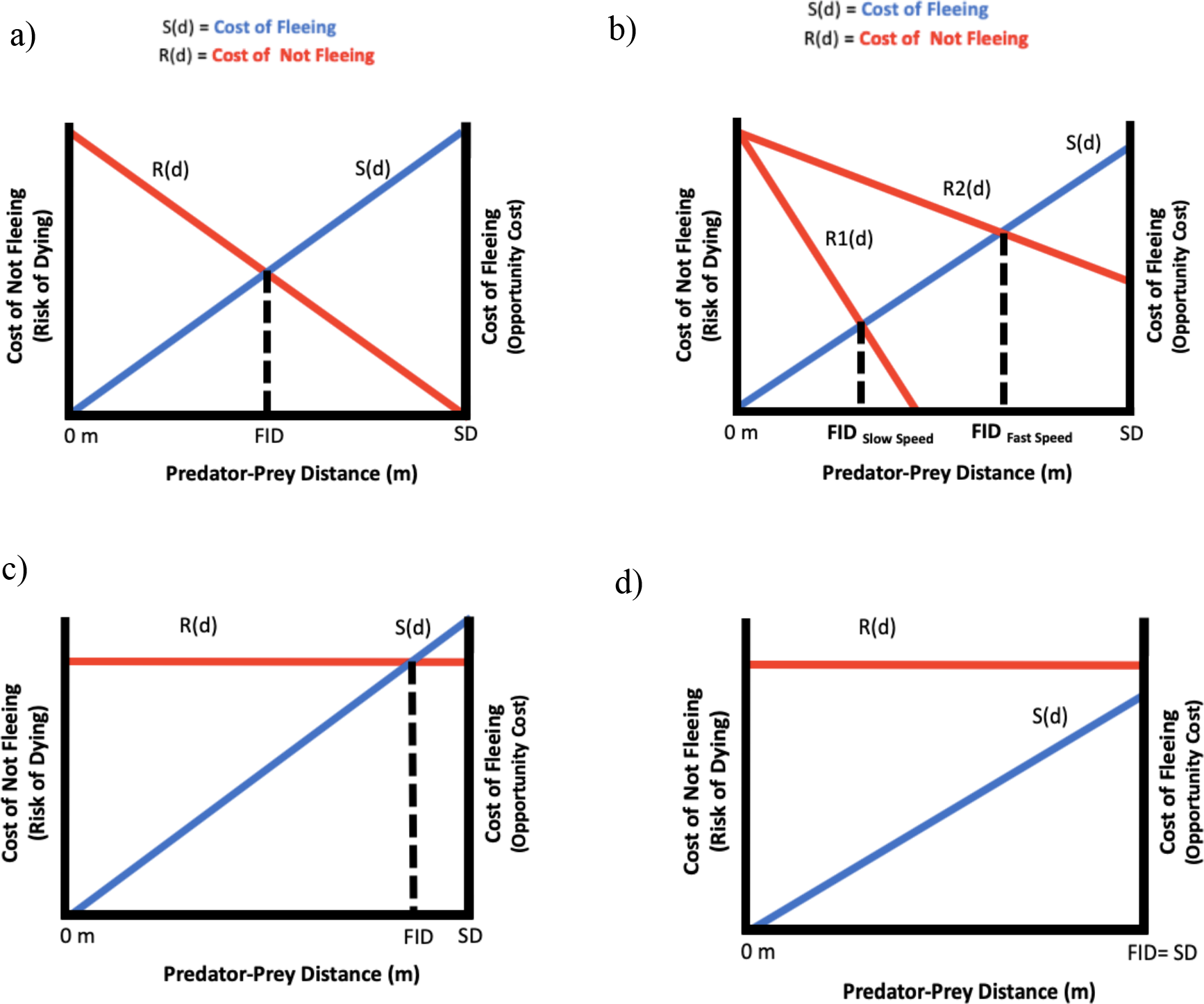
a) Economic escape model is a graphical model. The x-axis is the distance between an animal and an approaching threat (i.e., predator-prey distance). The red curve represents cost of not fleeing, R(d), and the blue curve represents the cost of fleeing, S(d). The x-coordinate where the two curves intersect is the predicted FID for the animal. b) As speed increases, the cost of not fleeing curve becomes more shallow and an animal’s predicted FID increases (R2(d)). As risk increases the absolute value of the slope for the cost of not fleeing curve approaches 0. c) If the two curves do intersect than FID becomes limited by the cost of fleeing. d) If the cost of fleeing is also shallow and the two cost curves do not intersect then FID = SD.

Ydenberg and Dill (1986) [18] portrayed the cost of fleeing curve as linear. The cost of not fleeing is associated with the risk of dying from the predator and its curve decreases with distance from the predator because the risk of dying decreases the farther away the animal is from the potential source of mortality (Fig 3a – red solid line). The model assumes that the intercept for the cost of not fleeing curve is the maximum possible loss of future fitness (i.e., dying), if the animal comes into contact with the predator (i.e., distance = 0) [25]. The original model portrayed the cost of not fleeing as nonlinear (i.e., exponential); however, the exact shape of the curve is unknown empirically, so we have represented it as linear (Fig 3a) for simplicity and following Cooper & Vitt (2002) [67]. The model predicts that the distance at which animals escape (i.e., FID) will be the intersection of the cost of not fleeing curve and the cost of fleeing curve because at this distance the reduction in future fitness will be minimized (Fig 3a – dotted black line). The exact relationship between FID and fitness is not specified; consequently, the shape of the curves are unknown [25, 67]. The original graphical model does not provide a mathematical formulation for either curve; however, Cooper and Frederick (2007) [25] suggested a mathematical formulation for an exponential or linear cost of not fleeing curves and a linear cost of fleeing curve (S1.1). Both cost curves are assumed to be monotonic [67].

The economic escape model has been the theoretical cornerstone for decades of escape behavior research [8]. While this graphical model maintains heuristic value, there are several gaps that should be addressed. First, the model depicts the cost of fleeing and the cost of not fleeing curve as independent of each other, yet it is possible that characteristics of the approaching predator simultaneously affect both cost curves. For example, if an animal is approached by a predator at a fast speed not only is there an increase in risk perception (an increase in the cost of not fleeing), but simultaneously there is less time to take advantage of a given opportunity (a decrease in the cost of fleeing). This potential lack of independence of the two curves can affect its predictions. Second, the curves should not be represented as equivalent. If an animal fails to escape then the consequence (i.e., death) results in a loss of all future fitness. In comparison, the one-time cost of fleeing prematurely from a predator is most likely non-consequential at that moment in time (i.e., less foraging time), and assuming that refugia present less risk. In that scenario, we would expect the slope of the cost of fleeing curve to be small (i.e., close to zero) if not constant. Third, the two curves impact future fitness on two different time scales. The cost of not escaping impacts future fitness immediately if a prey is captured by a predator, but the realized impact of the cost of fleeing on future fitness is the result of all cumulative opportunities the animal has left behind, and relative to resources elsewhere. Animals that tend to have a longer FID over the course of a lifetime would likely have a reduced future fitness compared to animals with a shorter FID. Careful consideration of the misalignment in temporal scales of the two costs curves is critical for the ability of this model to generate quantitative predictions (see below).

#### 2. Quantitative predictions

According to the economic escape model, the predicted FID will be at the intersection of the two cost curves. The relationship between the two curves should be either additive or multiplicative (S1.1). We can hypothetically explore the relationship between the two cost curves and fitness by either multiplying or adding the equations for the two cost curves put forth by Cooper and Frederick (2007) [25] for this model. This was done as a mathematical exercise and not based on any empirical data. We explored which relationship between the two curves, if any, produces a fitness curve that peaks for a single FID distance (S1.1). When the two cost curves are added and the absolute values of the slopes are equal, fitness does not change with the distance to the predator (S1.1), leading to no optimal FID regardless of the intersection in the two cost curves.

Alternatively, if we add the two cost curves and the absolute value of the slopes for the cost of not fleeing and the cost of fleeing are not equal, then the optimal FID is either 0 or FID is equal to the distance at which the predator initiates the approach (SD) (S1.1). Ideally, however, costs of remaining or fleeing should be expressed as likelihoods. Thus, only when the two curves are multiplied by each other is there a single peak in future fitness for a given predicted FID as a result of the two curves crossing (S1.1). The implication is that to generate quantitative predictions with this model the two cost curves should have a multiplicative relationship.

Unfortunately, the relationship between the cost curves and future fitness is unknown empirically. One possible set of proxies for the cost of not fleeing curve and the cost of fleeing is the probability of capture, or in the case of animal-vehicle interactions, the probability of collision with a vehicle and the probability of starvation based on FID [68, 69]. Survivorship is a critical factor in future fitness; so the probability of capture makes a suitable proxy for the cost of not fleeing curve [69]. The cost of fleeing might have a wide range of fitness consequences as a result of missed foraging or missed mating opportunities [18] . The economic escape model represents a single event and the interaction between an animal and an approaching vehicle is likely to happen on the order of seconds. Additionally, the decisions of animals to return to the spot where the interaction occurred can differ depending on whether the interaction was with a predator vs. a vehicle (i.e., shorter return times are expected with vehicles) or if the animal was missed by the vehicle or evaded (absent leaving the area) a collision without fitness-impacting effects. How these differences impact fitness is unclear empirically, which limits our ability to parameterize the model.

Overall, for the economic escape model to generate quantitative FID predictions we would need to parameterize both curves with empirical data. As discussed, generating a proxy for the cost of not fleeing curve is possible, but we have not found a suitable proxy for the cost of fleeing curve. Therefore, we could not generate quantitative predictions for this model in the context of animal-vehicle interactions.

#### 3. Sensitivity to approach speed

Because we could not generate quantitative predictions for the economic escape model, we did not explore its sensitivity to speed from a quantitative point of view.

### Blumstein’s economic escape model

#### 1. Overview

Blumstein’s economic escape model [24] attempts to provide an ultimate explanation because short-term escape decisions are linked to fitness. Blumstein’s model expands upon Ydenberg and Dill’s model [18] by partitioning the graphical model into three different zones, instead of only one, based on the response to the approaching predator (S1.2). The model starts at the distance where the predator begins its approach towards the animal (SD). Blumstein’s escape model assumes that the detection of the predator could occur in any of the three zones, and where detection occurs results in different behavioral outcomes. In other words, detection distance (DD)≤SD. In zone I, animals perceive a maximum amount of risk, and if they detect the predator in this zone, they escape immediately (Fig 4a). In zone II, upon detection animals monitor the approaching predator and escape at the distance that minimizes the reduction in fitness as determined by the intersection of the two cost curves (the cost of not fleeing and the cost of fleeing; Fig 4a), following from the economic escape model [18]. In zone III, animals do not escape because either they have detected the predator and perceive a minimum amount of risk or have not detected the predator at all (Fig 4a). Zones I and II and zones II and III are bounded by two threshold distances (the minimum distance, dmin; and the maximum distance, dmax; respectively) that establish where the animal will either escape immediately if a predator is detected closer than some minimum threshold distance or will not escape at all when a predator is beyond that maximum distance. Although the intersection of the cost curves determines the FID in zone II, as indicated for the economic escape model, animals escape immediately if the predator is detected closer than the predicted FID.

**Figure 4.**
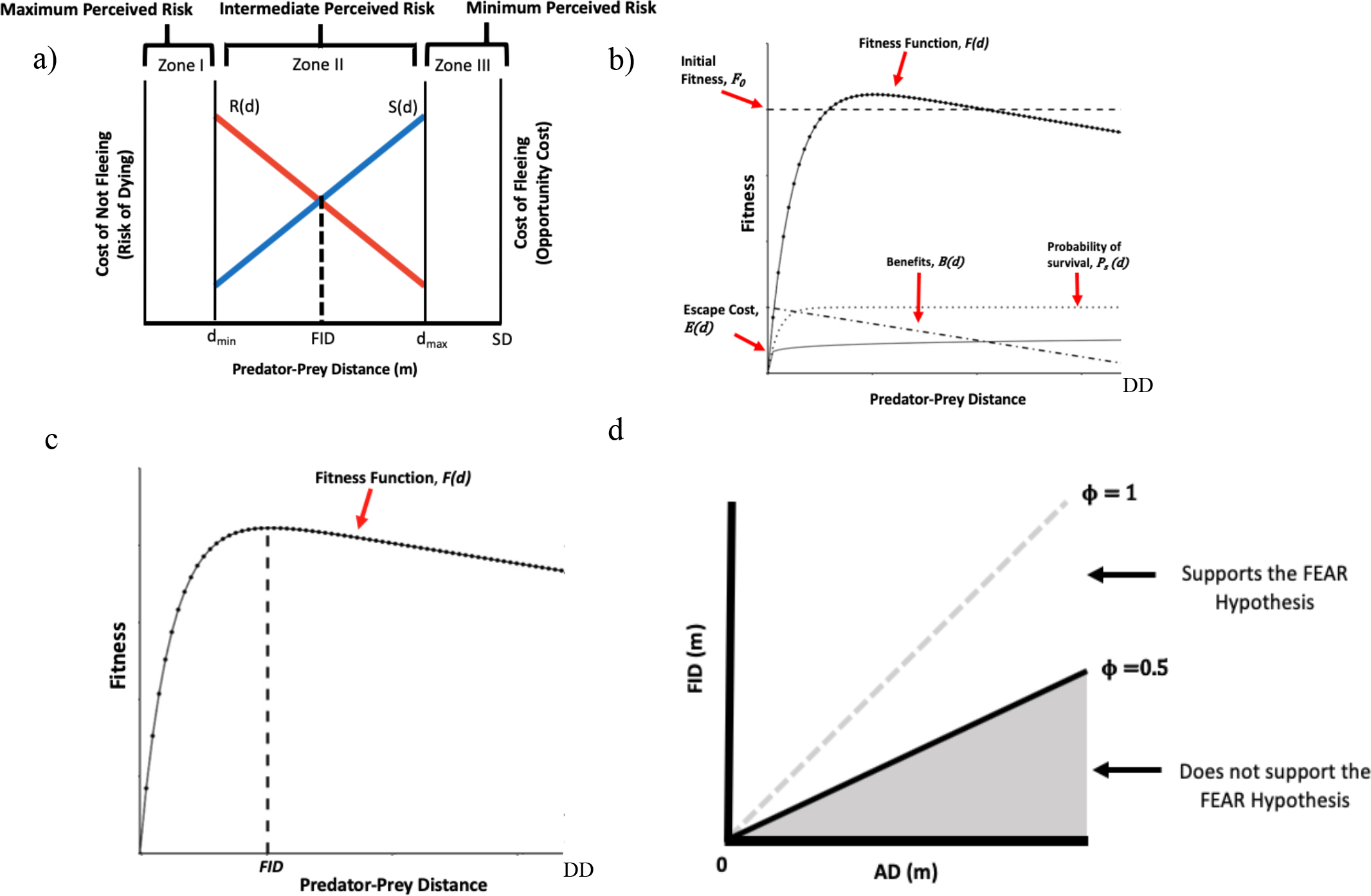
a) Graphical representation of Blumstein’s economic escape model. The x-axis is the distance between an immobile animal and an approaching threat (i.e., predator-prey distance). The red curve represents the cost of not fleeing, R(d), and the blue curve represents the cost of fleeing, S(d). a) In the original formulation Zone I, is from 0 to *dmin* and an animal always flee if a threat is detected. Zone II is from *dmin* to *dmax* and the predicted FID is determined by the animals assessment of the cost of not fleeing and the cost of fleeing. Zone II is similar to Economic escape model. Zone III is any distance greater than dmax and the animal does not respond to threats in this zone. b) Optimal escape model offers explicit mathematical formulations for an animal’s escape behavior. The x-axis is the distance between an immobile animal and an approaching threat. The interaction begins with the animal’s detection distance (DD). In optimal escape model an animal’s fitness, *F(d*), is determined by how initial Fitness ,(*F0*), the benefits gained during a threat’s approach, *B(d*), and the escape cost ,*E(d*), multiplied by the probability of survival, *Ps(d*), affect an animal’s initial fitness (*F0*). Fitness increases as the benefits functions increases at shorter distances away from the threat. Fitness then decreases as the probability of survival decreases at very close distances to the threat. Escape cost is a one-time cost the prey pays when it flees. Initial fitness does not change with distance. c) Animal escape (FID) at the peak of the fitness function (*F(d*)) in the optimal escape model. d) Illustration of the phi index was developed to determine if the AD vs. FID relationship supports the FEAR hypothesis. The range of ϕ is between 0 and 1. A ϕ value of 0 occurs when animals do not escape at all even after detection and a ϕ value of 1 occurs when prey escape the moment it detects the predator. ϕ must be greater than 0.5 to support the FEAR hypothesis (Samia & Blumstein 2014).

#### 2. Quantitative predictions

The logic of Blumstein’s economic escape model is similar to the economic escape model (see above). Zone II is limited to qualitative predictions because there is not empirical evidence on how the cost of fleeing is related to fitness, as discussed before. To generate quantitative predictions, the cost of fleeing and the cost of not fleeing curves should have a multiplicative relationship with each other, must be independent of each other, and have costs that can be comparable temporally. Consequently, it is not possible to generate quantitative FID predictions in the context of animal-vehicle interactions with this model.

#### 3. Sensitivity to approach speed

Because we could not generate quantitative predictions for the economic escape model, we did not explore its sensitivity to speed from a quantitative point of view.

### Optimal escape model

#### 1. Overview

The optimal escape model is different from the prior two models in that animals can enhance their fitness by optimizing the decision as to when to escape from a predator, as opposed to minimizing the cost to future fitness [25]. The optimal escape model can be considered an ultimate explanation for escape distance because escape decisions are expected to affect fitness. The model begins at the distance where the animal detects the approaching predator (DD). Future fitness is determined by how initial fitness is affected by three functions that vary relative to the distance from the predator: (1) benefits obtained from delaying escape (i.e., more time spent foraging instead of escaping), (2) the escape cost (i.e., the metabolic costs associated with escaping), and (3) the probability of survival (i.e., avoiding being caught as the predator approaches). Each function is made up of several parameters that generate a predicted FID (S1.3) [25]. An animal can enhance fitness by delaying escape (i.e., continue foraging) and allowing the predator to approach closer (Fig 4b). These benefits are offset by a one-time cost of escape, and ultimately weighted by the decreasing probability of survival as the predator gets closer (Fig 4b) [25]. Initial fitness remains constant regardless of the distance at which the animal decides to escape (S1.3). The model predicts that the animal will escape at the distance that optimizes fitness relative to all other distances (Fig 4c; S1.3) [25].

#### 2. Quantitative predictions

For the optimal escape model to make quantitative predictions about escape distance, empirical estimates of the fitness benefits obtained from delaying escape and the escape costs relative to initial fitness are necessary. However, we are unaware of any such empirical evidence over the span of a few seconds. Consequently, we could not generate quantitative predictions in the context of animal-vehicle collisions for the optimal escape model.

#### 3. Sensitivity to approach speed

Because we could not generate quantitative predictions for the economic escape model, we did not explore its sensitivity to speed from a quantitative point of view.

### Perceptual limits hypothesis

#### 1. Overview

The perceptual limits hypothesis is a proximate explanation for escape behavior because it states that escape is limited by the perceptual abilities of the animal (i.e., the ability of its sensory system to resolve the approaching predator) [18, 26]. The perceptual limits hypothesis posits that animals should escape immediately after they detect a predator. Therefore, this hypothesis predicts that flight initiation distance will be the same as detection distance (FID = DD, S1.4).

From the perspective of a prey, the interaction begins once it has detected the predator, but this is not stated explicitly. The perceptual limits hypothesis establishes that animals do not assess an approaching predator because escape occurs immediately after detection. The difference between the predator’s starting distance (SD) and the prey’s detection distance (DD) is likely to be associated with differences in sensory constraints between individuals or between species (S1.4). Consequently, individuals within a species that differ in their sensory resolution [70, 71] are predicted to have different escape distances. Similarly, differences between species in sensory resolution are [72] predicted to result in different FIDs. Individuals/species with a more constrained sensory system are expected to detect predators at closer distances, and to have shorter FIDs relative to individuals/species with less constrained sensory systems (S1.4). The implication is that the distance at which animals escape will be a function of the constraints of the sensory dimension/s used to detect predators.

#### 2. Quantitative predictions

The perceptual limits hypothesis does not have any mathematical or graphical framework to generate predictions. However, given its reliance on the limits of the sensory system to resolve the stimulus (i.e., a predator) relative to the background and the fact that birds are visually oriented organisms, we followed Tyrrell & Fernandez-Juricic (2015) [73] and considered visual acuity as a proxy to estimate DD. Visual acuity is the ability of an organism to resolve two objects as different at a maximum theoretically possible distance. Visual acuity, measured in cycles per degree (i.e., one cycle consists of one light bar plus one dark bar of a grating per degree subtended at the eye), can be estimated anatomically using measurements of eye size (axial length) and retinal cell density (e.g., photoreceptor cells or retinal ganglion cell densities) [38, 74] .

If we know the size of the approaching vehicle and the visual acuity for a given individual/species, we can estimate the distance over which the object could be resolved by the eyes [74, 75]. There are some assumptions associated with this calculation: (1) the object occupies one degree of a species visual field at the detection distance, (2) the animal will be able to identify the threat at the moment of detection, (3) the animal views the approaching threat monocularly, and (4) viewing occurs under optimal ambient light conditions (S1.4) [75, 76].

Assumptions notwithstanding, this estimated distance can be considered a proxy of the distance at which an animal could detect a threat, and according to the perceptual limits hypothesis, this is also the FID of that species/individual.

To exemplify the ability of the perceptual limit hypothesis to make quantitative predictions based on the data from DeVault et al. (2015) [15], we used the visual acuity of brown- headed cowbirds (4.82 cycles per degree) [38]. We estimated that it would detect an approaching vehicle (assumed width of a car = 1.73 meters) [77] 475 m away (S1.4). Since DD is equal to FID, the perceptual limit hypothesis predicts that brown-headed cowbirds would show a FID distance of 475 m in response to an approaching vehicle.

#### 3. Sensitivity to approach speed

Despite the fact that the model allowed us to generate quantitative predictions, we could not assess its sensitivity to speed because the perceptual limit hypothesis does not incorporate any parameter related to speed. Detection in the formulation used to generate quantitative predictions is a function of distance away from the viewer, irrespective of its speed.

### Flush early and avoid the rush (FEAR) hypothesis

#### 1. Overview

The flush early and avoid the rush (FEAR) hypothesis offers an ultimate explanation of escape behavior because the distance at which an animal escapes is based on short-term decisions, including how the animal allocates monitoring cost, which will ultimately affect fitness. The FEAR hypothesis argues that animals tend to escape early to minimize the cost of monitoring an approaching predator because they have limited attention: monitoring the predator diverts attention away from other potential fitness enhancing activities, such as foraging, mating, etc. [8,27,51–53] (S1.5). The interaction begins at the point where the predator starts the approach towards the animal (SD). The FEAR hypothesis assumes that animals detect an approaching predator at the distance where they become alert, such that DD = AD (Blumstein 2010; S1.5).

Animals begin to incur the monitoring costs once they have detected the approaching predator. AD is assumed to be greater than or equal to FID (Blumstein 2010; S1.5). Consequently, the possible range for FID values is constrained in the following way: 0 *<* FID *≤* AD *≤* SD (S1.5).

Higher attentional costs are associated with longer monitoring times [27, 40]. After detecting a predator that moves at a constant speed, the more the animal delays its escape, the higher its monitoring costs become because of the increased monitoring duration preventing the animal from directing its attention towards other fitness enhancing activities [27, 78]. Although the animal should escape after detecting the predator to reduce the cost associated with monitoring, it does not escape immediately after detection because it assesses the predator approaching. Thus, the predicted FID is the result of a decision between the cost of fleeing and the cost of not fleeing, like the economic escape model [78]. If the animal detects the predator farther away (long AD), the distance at which they escape is also expected to be farther away (long FID) to reduce monitoring costs. The FEAR hypothesis therefore predicts that AD and FID should be positively correlated (S1.5) [27,79,80].

#### 2. Quantitative predictions

In the FEAR hypothesis, AD is a predictor of FID and by definition AD must be greater than or equal to FID, which causes statistical complications when testing this prediction quantitatively. For instance, Dumont et al. (2012) [81] showed that the FEAR hypothesis leads to statistically significant and positive relationships using random data. This is because as the values of AD increase, the range of possible FID values from a random uniform distribution also increases, which will violates the homogeneity of variance assumption in a linear model [81]. This scenario can result in AD being a spurious significant predictor of FID [81]. Three solutions have been proposed to overcome this issue.

The first proposed solution is to simulate AD and FID values from a uniform distribution, maintaining the same aforementioned constraints (0 *<* FID *≤* AD *≤* SD) and then compare the simulated AD vs. FID slopes with the empirically observed AD vs. FID slope [81]. If the empirically observed slope falls outside of the 95% confidence interval of the simulated slopes, the observed AD vs. FID relationship would then support the FEAR hypothesis rather than being a mathematical artefact [81]. However, outliers can over- or under-estimate the value of the slope [82]. The second proposed solution is to use quantile regression, which essentially attempts to fit a linear trend on a particular portion of the data. Chamaillé-Jammes & Blumstein [83] proposed to analyze the 10% lowest quantile of the observed AD vs. FID data to minimize the homogeneity of variance issue. Although quantile regression is robust to outliers, this approach is only accurate with larger sample sizes, which may not be easy to obtain [81,84,85]. The third proposed solution is to use the phi-index, *Φ*, which is the standardized distance between the expected AD and observed FID in an AD vs. FID space where the intercept is 0 [42]. The phi-index estimates how close the observed relationship between AD vs. FID is from a 1:1 relationship [42]. The phi- index can be considered a proxy for the effect size of AD on FID. Mathematically, *Φ* can vary between 0 (i.e., animals do not escape at all even after detection) and 1 (i.e., animals escape the moment they detect the predator) [42]. In the AD vs. FID space, the greater the numerical difference between AD and FID, the lower the *Φ* value, suggesting that animals delay escape in a way that is not consistent with the FEAR hypothesis. The smaller the numerical difference between AD and FID (Fig 4d), the higher the *Φ* value, which is consistent with the FEAR hypothesis (i.e., animals escape after detecting a predator). The null expectation for the *Φ* index is 0.5 [42] at which point animal responses are exactly halfway between escaping after detection and not escaping at all. Therefore, the *Φ* value is expected to be greater than 0.5 to support the FEAR hypothesis.

To establish whether the observed *Φ* value is significantly greater than 0.5, 1,000 *Φ* values are simulated from a random FID uniform distribution for each empirically observed AD, where FID *≤* AD [42]. The 1,000 *Φ* values are compared to the empirically observed *Φ* value. If the observed *Φ* value is greater than at least 5% of the simulated values, then we can reject the null hypothesis that AD and FID vary randomly [42] . An observed *Φ* value only supports the FEAR hypothesis when it is both greater than 0.5 and significant (p-value < 0.05). The phi index is preferred over the previous two methods as it is not sensitive to outliers, nor does it require a large sample size (S1.5) [8,42,82].

To generate quantitative predictions according to the FEAR hypothesis with the DeVault et al. (2015) dataset (n=120), we fitted the equation of the line in the AD vs. FID space. The intercept of an AD vs. FID relationship was assumed to be 0 because the hypothesis assumes that detection precedes escape; therefore, if AD is equal to 0, then FID should be equal to 0 (S1.5) [42]. We decided not to use *Φ* as a proxy for the slope because of its tendency to overestimate the value of the AD vs FID slope compared to a linear model (S1.5), and instead used the observed slope of the AD vs. FID relationship, which ranges from 0 to 1. The maximum possible slope is 1, because FID is constrained by AD (i.e., FID *≤* AD); thus, an animal’s longest possible FID occurs when the animal escapes immediately upon detection (i.e., FID = AD) [42]. The minimum possible slope of an AD vs FID relationship is 0 because of the same constraints on FID relative to AD [27,42,78] . We generated predictions on FID by multiplying the observed slope of the linear regression by the empirically observed mean AD at the 95th percentile [86–88] for each speed treatment for a total of eight different speeds.

We first estimated *Φ* for the observed AD and FIDs for each of the eight speed treatments in DeVault et al. (2015) [15]. Then, we assessed if each *Φ* value was significantly different from 0.5. We found that for every speed treatment *Φ* was significantly greater than 0.5 thus each speed treatment was consistent with the FEAR hypothesis. We then ran linear models on the AD vs. FID relationship for each speed treatment, and obtained the predicted FID for each speed by multiplying the model predicted slope by the AD at the 95th percentile. We were able to generate quantitative FID predictions using the FEAR hypothesis, with FID varying from 9.12 to 19.59 m across different speeds with a mean of 14.53.

#### 3. Sensitivity to speed

The relationship between effect size (*Φ*) and the slope of the FID vs. approach speed relationship resembled a sigmoidal curve (Fig 5a). *Φ* values greater than 0.5 suggest that escape behavior follows the FEAR hypothesis [42]. All negative slope values had a mean *Φ* less than 0.5 (mean 0.025, SD = 0.044), suggesting that in that range of the FID vs. approach speed relationship, the FEAR hypothesis is not sensitive to approach speed. When the slope value was in the range of slopes not significantly different from 0, we found that *Φ* was higher than 0.50 (mean 0.703, SD = 0.033), suggesting that the FEAR hypothesis is sensitive to approach speed in that range of the FID vs. approach speed relationship. For all positive slope values, *Φ* was higher than that 0.5 (mean 0.993, SD = 0.012), suggesting that the FEAR hypothesis is sensitive to approach speed in that range of the FID vs. approach speed relationship.

**Figure 5.**
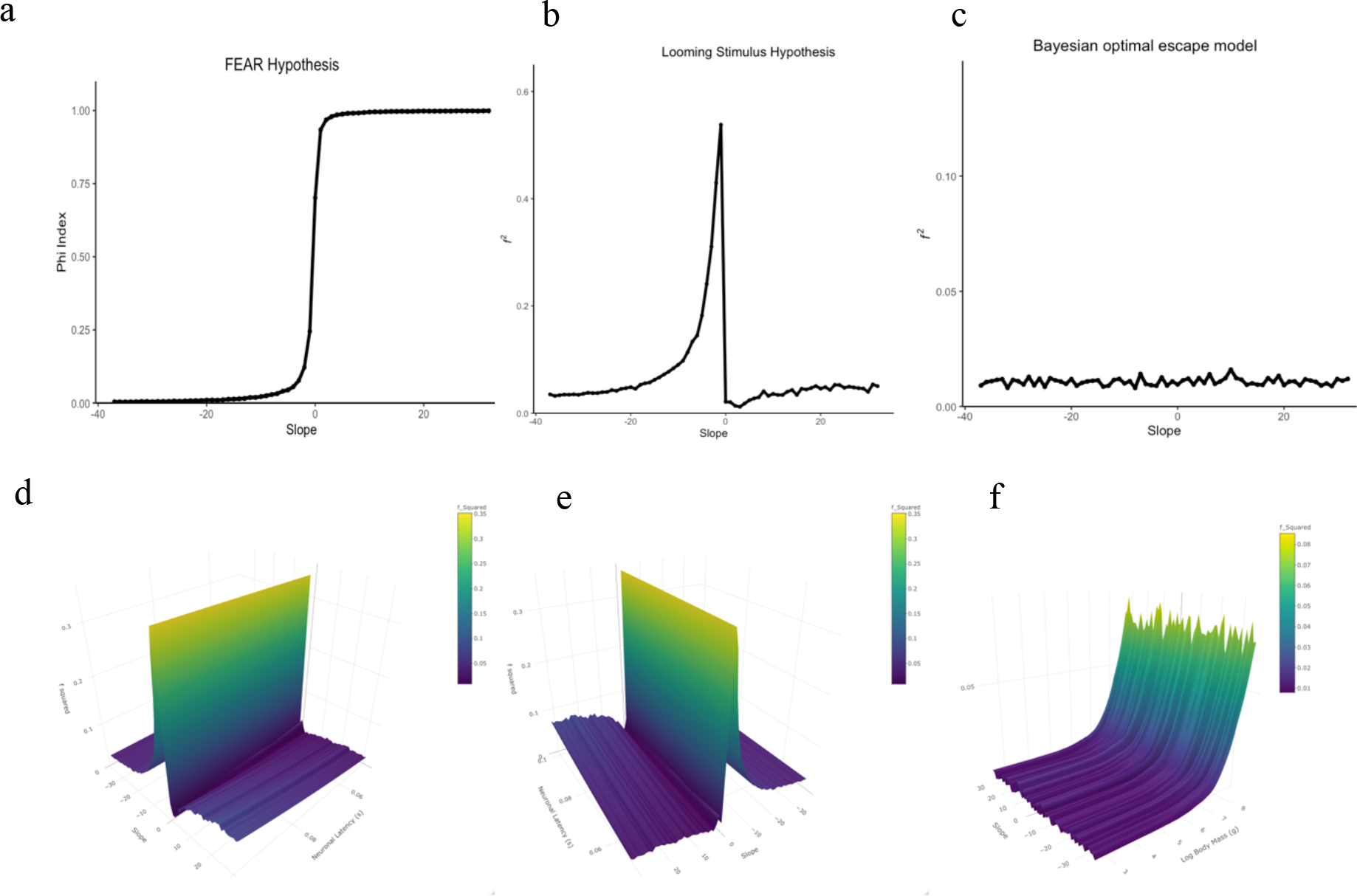
a) The mean phi-index, the effect size metric for the FEAR hypothesis, across iterations for each slope. The figure suggest that model is most sensitive to vehicle approach speed when the slope is equal to or greater than zero. b) The mean *f ^2^* across iterations for each slope according to the looming hypothesis when latency is 0.075 seconds, which suggest the model is sensitive to the delayed margin of safety. c) The mean *f ^2^* across iterations for each slope according to the Bayesian optimal escape model when the body mass is 267.4 grams, which suggest the model is not sensitive to approach sped. d & e) The mean *f ^2^* across iterations for each combination of slope and neuronal latency value for the looming stimulus hypothesis. d and e show the same graph but from two different viewpoints. Across neuronal latency values it seems the model is sensitive to the speed effect at slightly negative slopes. f) The mean *f ^2^* across iterations for each combination of slope and body mass value for the Bayesian optimal escape model. The figure suggests that only at a size of larger then 1kg is the model FID predictions sensitive to vehicle approach speed.

### Looming stimulus hypothesis

#### 1. Overview

The looming stimulus hypothesis provides a proximate explanation for escape behavior because the decision to escape is triggered by the reaction of certain neurons to an approaching threat. A looming stimulus represents an object (predator, vehicle) moving along an impending collision path towards the viewer [28]. If we assume that the approaching stimulus is a 2D object, the size of the stimulus can be measured from the perspective of the viewer’s visual angle. Visual angle is the angle (*θ* degrees) subtended by the approaching object onto the retina where the size of the angle is proportional to the size of the stimulus [29]. As the object approaches closer, the visual angle expands. The faster the object approaches, the rate of visual angle expansion increases (*θ’* degrees per second). Both *θ* and *θ’* are a function of the remaining time before the object collides with the viewer, also known as time to collision (TTC). If the speed of the approaching object is known, TTC can be converted into the distance between the object and the animal (S1.6).

The looming stimulus hypothesis states that animals escape because of neurons firing when the ratio of the visual angle, *θ* (t), and the rate at which the visual angle expands, *θ* (t), referred to as the optical variable tau (*τ*), decreases to a critical threshold (S1.6) [28,29,89]. Tau is essentially time-to-collision if approach velocity is constant. *τ*decreases because the change in *θ’* (t) as time to collision decreases is greater than the change in *θ* (t) (S1.6). At some threshold value of *τ*, neurons begin firing, and after a certain neuronal latency (*δ*), they reach their peak firing rate, triggering the escape response (S1.6). The hypothesis does not specify a threshold value of *τ* at which neurons activate. In pigeons, neurons characterized as encoding *τ* begin firing between 0.3 sec to 1.4 sec relative to objects of varying speeds and sizes, where neuronal latency (*δ*) between the onset of neuron firing and the peak firing rate was 0.5 to 1 second [90–92]. A proxy for *τ* is the ratio of object size to object approach speed [93]. An escape response is dependent on the animal continually receiving visual information about the looming object; therefore, the looming hypothesis assumes that the animal has detected and continually monitors the approaching object.

#### 2. Quantitative predictions

The looming stimulus hypothesis can generate quantitative FID predictions with a threshold value of *τ* at which neurons begin to respond to an approaching object. If animals escape at some threshold value of *τ* and if the object size and approach speed is known, assuming approach speed is constant, FID can be estimated from the visual angle at which the approaching object reaches the critical value of *τ* (Gabbiani, personal communication). However, to estimate the threshold value *τ* from behavioral data requires the assumption that at the moment of escape (i.e., FID), the neurons encoding *τ* reach peak firing, generating the response [66,90,92].

According to the looming stimulus hypothesis, FID is a function of the threshold value of *τ* and the neuronal latency exhibited by that species (*δ*). Without an a priori estimate of the threshold value of *τ*, we estimated the effect of neuronal latency (*δ*) on FID at different approach speeds using the DeVault et al. (2015) [15]. dataset. Neuronal latency can be estimated with a linear regression between the ratio of vehicle size to approach speed, a proxy for *τ*, as the independent variable and the average TTC*Flight* as the dependent variable (following Fotowat & Gabbiani 2011[89]). We considered the intercept of that model as a proxy of *δ* (following Fotowat & Gabbiani 2011[89]). The estimated *δ* was 0.0962 seconds (S1.6). According to the model, the predicted FID was estimated as: 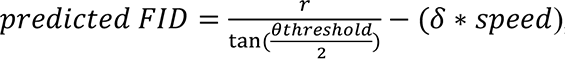, where *r* is the radius of the approaching object and *θ threshold* is the visual angle at which the animal fled (i.e., FID). The part of the equation 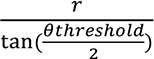 is the trigonometric definition of FID. We substituted the average FID at different approach speeds (using DeVault et al. 2015 data [15]) for the predicted FID estimated from the threshold value *τ*. The neuronal latency was subtracted from average FID at different approach speeds because a delay in escape results in a shorter FID. We multiplied *δ* by approach speed to convert the delay from time units into distance units. We were able to generate quantitative predictions using the looming stimulus hypothesis, with FID varying from 24.53 to 33.23 m (mean = 30.03 m) across different speeds. However, we acknowledge that without an estimate of the threshold value of *τ* our approach of generating quantitative prediction is limited. We followed this approach though because it is useful to understand how neuronal latency impacts FID at different approach speeds.

#### 3. Sensitivity to speed

In the looming stimulus hypothesis, the relationship between effect size (*f^2^)* and the slope of the FID vs. approach speed relationship increased from -37 to -1 exponentially, then dropped substantially in the range of slopes not significantly different from 0, eventually increasing and then leveling off as the slope reached 32 (Fig 5b). Across all negative slopes, *f^2^* varied between small and medium (mean 0.096, SD= 0.112). *f^2^* was closer to a small effect size when the slopes ranged from -37 to -11. From a slope of -10 to a slope of -1, *f^2^* began to increase beyond a medium effect size and eventually reaching a large effect size. Across all negative slopes, the effect sizes suggests that the looming stimulus hypothesis is sensitive to approach speed. In the range of slopes not significantly different from 0, *f^2^* was less than a small effect size anticipated by Cohen 1992[41] (mean 0.021), suggesting that the looming stimulus hypothesis is not sensitive to approach speed. Across all positive slopes, *f^2^* was small (mean 0.039, SD = 0.012), suggesting that the looming stimulus hypothesis is most likely not sensitive to approach speed. These relationships between effect size and slope had a fixed neuronal latency (see Methods).

When we allowed neuronal latency to vary, effect size changed with slope at different neuronal latency values (Figs 5d, 5e). Across different neuronal latency values, the curve is relatively qualitatively similar to Fig 5b across the range of slopes (-37 to 32). As slope increases from -37 to -1 (Figs 5d, 5e), different neuronal latency values do not result in noticeable differences in *f ^2^* relative to Fig 5b. In the range of slopes not significantly different from 0, *f^2^* still dropped substantially, but tended to be smaller for shorter neuronal latency values compared to longer neuronal latency values (Figs 5d, 5e). From a slope of 1 to 32, *f^2^* also slowly increased and then leveled off (Figs 5d, 5e). Although longer neuronal latency values tended to yield larger *f^2^*, effect sizes still remained small (Figs 5d, 5e). These patterns suggests that, irrespective of neuronal latency, the looming stimulus hypothesis is sensitive to approach speed for negative slopes (-10 to -1).

### Visual cue model

#### 1. Overview

The visual cue model [30] offers a proximate mechanism to account for escape behavior as the escape decision is based on the distinct change in the perceived size (i.e., visual angle) of an object approaching; similar to the looming stimulus hypothesis. The model assumes that animals have detected the approaching object to make predictions about escape behavior. The model uses a proxy for the concept of *τ* introduced in the looming stimulus hypothesis section referred to as *A*, which is estimated as the ratio of the approaching object perceived profile size relative to distance from the predator. The implication is that in the visual cue model, the speed of the approaching object is constant and not explicitly considered like in the looming stimulus hypothesis. Profile size is defined as the diameter of the approaching object multiplied by a shape-specific coefficient *κ* [30]. The visual cue model argues that animals rely on changes in the *A* at two different distances, *ΔA*. When *ΔA* exceeds some threshold, animals escape. This is another difference with the looming stimulus hypothesis, which considers the continuous expansion of the stimulus in the visual field of the observer during the approach.

The model incorporates other parameters, such as the approaching object trajectory (i.e., direct vs. tangential approach), vegetation structure (i.e., varying from 0 – low habitat complexity – to 1 – high habitat complexity–), and individual differences (i.e., indicating the variation between individuals around the population mean in escape responses) [30]. For instance, a tangential approach tends to lead to shorter FIDs than a direct approach; more complex habitat structure tends to lead to shorter FIDs than less complex habitat structure; etc. (S1.7) [30].

#### 2. Quantitative predictions

We generated quantitative predictions with the visual cue model following the procedures outlined in Javurkova et al. (2012) [30]. We set the model equal to zero and solved for FID using the data from DeVault et al. (2015) [15] and following Javurkova et al. 2012 [30] (S1.7). We had to make additional assumptions. First, we assumed that the vehicle approached the animal directly (i.e., approach angle = 0 degrees) to be consistent with the other models we studied. Second, we assumed there was no vegetation blocking the path of the vehicle approaching (i.e., vegetation = 0) (S1.7). Third, we assumed that there were no behavioral differences for the sake of simplicity. Fourth, we assumed that the profile size for an approaching vehicle was circular with a diameter of 1.73 m (S1.7) [15,30,77] .

The visual cue model requires establishing two different distances at which the animal sees the threat. We estimated the first distance at 475 m because that is the maximum theoretically possible detection distance based on a brown-headed cowbird visual acuity [38, 75]. We used the mean AD (from DeVault et al. 2015 [15]) as a proxy for the second distance at which a brown-headed cowbird might assess the approaching vehicle. We estimated the threshold value of *ΔA*, the change in the ratio of profile size relative to distance at which an animal escapes, based on the mean TTC*flight* for each speed treatment in DeVault et al. (2015) [15]. We used the NLRoot package in R to estimate the predicted FID with the bisection method as described by Javurkova et al. 2012 [30] (S1.7) [94]. We were able to generate quantitative FID estimates with the visual cue model, with FID varying from 16.10 to 21.9 m across different speeds. The average predicted FID across speed treatments was 19.25 m.

#### 3. Sensitivity to speed

Although we generated quantitative predictions, we could not assess the sensitivity to speed of the visual cue model because it does not incorporate any parameter related directly (or indirectly) to speed. The perceived change in size when viewed at the two different distances proposed by the model for an object moving at different speeds will be the same.

### Bayesian optimal escape model

#### 1. Overview

The Bayesian optimal escape model offers an ultimate explanation for escape behavior because animals evaluate the energetic cost of escaping relative to the energetic cost of remaining to determine the optimal FID, and both costs are assumed to be related to fitness [31]. The model starts at the point where animals detect the predator, which is assumed to be the AD. In the Bayesian optimal escape model, animals escape (i.e., FID) at the point where the energetic cost of fleeing is equal to or less than the perceived energetic cost of remaining, which is weighted by their perceived probability of attack [31]. The energetic cost of fleeing is the metabolic cost associated with escaping from an approaching predator. The perceived energetic cost of remaining is based on the current energetic reserves or how much energy it can afford to lose in an encounter with a predator (e.g., daily energy expenditure) [31].

The perceived probability of attack is estimated with Bayes theorem, and it is a function of the behavior of the approaching predator. More specifically, the perceived probability of attack depends on two functions: the proportional distance to the predator after detection, and the proportion of the predator approach speed relative to the prey’s maximum escape speed [31]. The model also incorporates a coefficient (*α*) reflecting the behavioral bias of the animal (past experience, habituation, or internal state) [31]. The coefficient *α* varies between 0 (i.e., an animal tends to have a shorter FID) and 1 (i.e., an animal tends to have a longer FID).

#### 2. Quantitative predictions

We were able to generate quantitative predictions with this model using the data from DeVault et al. (2015). We used the risk functions provided by Sutton & O’Dwyer (2018) [31] (S1.8), although alternative risk functions could be implemented. We parameterized the model with the observed cowbird escape speed (3.75 m/s) and mean AD for each speed treatment [15]. We assumed that the vehicle approached the cowbird directly (i.e., approach angle = 0°) to make the model’s predictions comparable with previous models. We did not have empirical information to estimate the coefficient of behavioral bias, *α*; therefore, we assumed no bias (*α* = 0.5) [31].

The metabolic cost of fleeing from an approaching vehicle was estimated as: W = 61.72*Mb^0.79^*, where *Mb* is body mass (kg), to estimate the metabolic cost of short flight in passerines [35]. Brown-headed cowbirds needed 0.8 sec to travel 3 m and avoid a collision [15]; thus, we assumed the metabolic cost of escape was 7.83 x 10^-3^ kJ. We estimated the brown-headed cowbird daily energy expenditure (kJ) based on their mean body mass (43.9 g; female body mass = 38.8 g, male body mass = 49 g) [36]. We used Daan’s et al. 1990 [34] equation to estimate daily energy expenditure: DEE = -0.80 + 0.66*M* (S1.8). With the Bayesian optimal escape model, we predicted that FID would vary from 38.92 to 48.98 m across different speeds. The average predicted FID across speed treatments was 44.99 m.

#### 3. Sensitivity to speed

In the Bayesian optimal escape model, the relationship between effect size (*f^2^)* and the slope of the FID vs. approach speed relationship did not vary much regardless of whether the slope ranged from negative, to zero, to positive (Fig 5c). Across all negative slopes values (mean 0.011, SD = 0.001), in the range of slopes was close to 0 (mean 0.009), and across all positive slopes values (mean 0.011, SD = 0.002), *f^2^* had less than a small effect size according to Cohen 1992 [41] (Fig 5c), suggesting that the Bayesian optimal escape model is not sensitive to approach speed. These relationships between effect size and slope had a fixed body mass (see Methods).

When we allowed body mass to vary (Fig 5f), the relationship between effect size and slope varied with different body mass values. For most body mass values, *f^2^* was less than a small effect size (Fig 5f), following the patterns presented in Fig 5c. However, when body mass was greater than 1.1 kg (log body mass of 7 grams in Fig 5f), *f^2^* began to increase exponentially reaching a medium effect size at the largest observed body mass of 6.25 kg. Throughout these higher body masses, effect size did not vary substantially with slope (Fig 5f). Overall, the Bayesian optimal escape model appears sensitive to vehicle approach speed for species with large body masses.

## Discussion

Our findings are summarized in Table 1. Five models provide an ultimate explanation (i.e., Economic Escape Model, Blumstein’s Economic Escape Model, optimal escape model, FEAR hypothesis, Bayesian optimal escape model) and three models provide a proximate explanation (i.e., perceptual limits hypothesis, looming stimulus hypothesis, visual cue model) for the distance at which animal are expected to escape from an approaching threat (predator, vehicle, etc.).

**Table 1.** We characterized each model based on its ability to generate either qualitative or quantitative predictions with different approach speeds. When possible we generated quantitative FID predictions with the model. We identified whether the model explicitly includes a parameter relative to the threats approach speed. Lastly we determined whether the model was sensitive to vehicle approach speed based on figure 5.

| Model name | Can the model generate qualitative predictions relative to speed? | Can the model generate quantitative predictions relative to speed? | Quantitative FID predictions | Does the model have a parameter for speed? | Sensitive to vehicle approach speed |
| --- | --- | --- | --- | --- | --- |
| Economic escape model | Yes | No | N/A | No | N/A |
| Blumstein's economic escape model | Yes | No | N/A | No | N/A |
| Optimal escape model | Yes | No | N/A | No | N/A |
| The perceptual limits hypothesis | No | No | FID = 474.67 m | No | No |
| Flush early and avoid the rush (FEAR) hypothesis | Yes | Yes, implicitly | mean FID = 14.53 m | No | Yes, spatial & temporal margin of safety only |
| The looming stimulus hypothesis | Yes | Yes | mean FID = 30.03 m | Yes | Yes, speed effect only |
| Visual cue model | No | No | mean FID = 19.25 | No | No |
| Bayesian optimal escape model | Yes | Yes | mean FID = 44.99 m | Yes | Yes, for spatial, temporal margin of safety, speed effect for species > 1kg. |

All models with the exception of the perceptual limits hypothesis and the visual cue model (Table 1) could generate *qualitative* predictions about changes in FID (i.e., longer FID, shorter FID) with changes in the speed of the object approaching (i.e., high speed, low speed). In fact, some of these models were used in making *qualitative* predictions in empirical studies with birds [14, 15]. Qualitative predictions certainly have value in the initial investigation of how animals react to high-speed vehicles; yet, the ability to generate *quantitative* predictions about escape distance can provide an opportunity to distinguish between predictions from different models, improving our ability to establish the degree to which responses from different species vary in different environments, and consequently enhance our capacity to design and implement management practices that could minimize animal-vehicle collisions.

Five of the eight models were able to generate *quantitative* predictions about escape distances of a high-speed vehicle (Table 1). The main reason three of the models could not generate quantitative predictions was because the range of values of some of their key parameters are unknown at this time. Future research into these parameters, such as the cost of fleeing in terms of fitness, could unlock the potential of these models to predict quantitatively escape distances.

When comparing the mean predicted FID across models (Table 1), the values varied from m to 475 m. The order of magnitude differences between the perceptual limits hypothesis and the other four models (FEAR hypothesis, looming stimulus hypothesis, visual cue model, Bayesian optimal escape model; Table 1) may be related to their very different assumptions. In the perceptual limits hypothesis, escape (FID) occurs at the moment of detection (DD); however, in the other four models, escape occurs *after* detection (Fig 6), and our knowledge of species-specific points of detection is limited. This difference likely leads to an overestimation of escape distance in the perceptual limits hypothesis relative to the other four models (Table 1). Additionally, the looming stimulus hypothesis and the visual cue model had similar mechanism (i.e., visual angle increases as a threat approaches) as well as their similar assumption by which DD is higher than FID, whereas no AD is explicitly considered in the model formulation (Fig 6). Finally, the FEAR hypothesis and the Bayesian optimal escape model both consider that AD is a proxy of DD, and that AD is higher than FID (Fig 6). The assumption that animals become alert at the moment of detection is a useful simplification for modeling purposes, but the evidence in humans contradicts it because visual detection of an object tends to occur considerably before visual inspection [95–97] , and potentially alert behavior. The bottom line is that the studied models differ to a large extent in their assumptions about the timing of detection, alert, and escape. This could limit our ability to apply certain models depending on the biology of the study organism.

**Figure 6.**
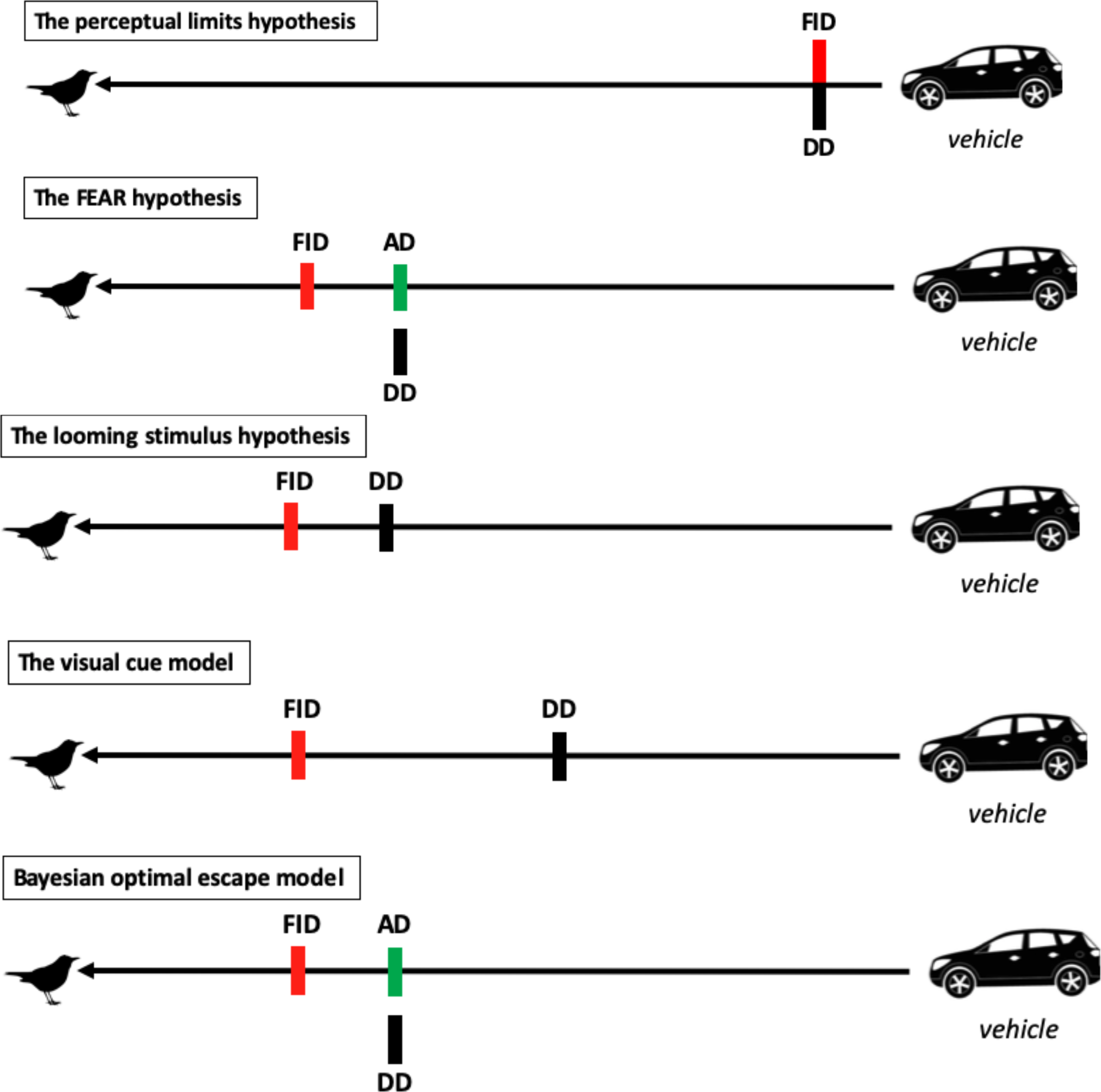
The illustrated differences in the detection assumptions for the five models capable of producing quantitative predictions. FID refers to flight initiation distance, AD refers to alert distances, and DD refers to detection distance.

Of the five models that allow quantitative predictions, we were able to assess their sensitivity to speed for only three of them: the FEAR hypothesis, the looming stimulus hypothesis, and the Bayesian optimal escape model. The looming stimulus hypothesis and the Bayesian optimal escape model both explicitly incorporate a parameter establishing the approach speed of the threat . While the FEAR hypothesis does not explicitly incorporate a parameter to account for approach speed [38, 98] , we were able to indirectly evaluate the effects of speed by considering how approach speed affects an AD and FID relationship with the phi index [42].

Our simulation results suggest that sensitivity to approach speed (i.e., how much vehicle speeds accounts for the variation in FID) is a function of the model and the rules of escape animals use (temporal margin of safety, spatial margin of safety, delayed margin of safety). The FEAR hypothesis is sensitive to vehicle approach speeds only when an animal follows the temporal or spatial margin of safety; the looming stimulus hypothesis is sensitive to vehicle approach speeds only when an animal follows the delayed margin of safety; and the Bayesian optimal escape model becomes sensitive to approach speed, regardless of the escape rule, only for species larger than 1 kg. Therefore, generating FID predictions relative to vehicle approach speed should be highly dependent on establishing the rules of escape of the target species in response to high-speed vehicles [14, 15] . Overall, we suggest that using models of escape behavior developed for predator-prey interactions to generate quantitative predictions about responses of animals to high-speed approaching vehicles should be applied with much caution.

Additionally, applying models to a target species should involve considering to what degree model assumptions are met. For instance, whether or not a species detects the approaching vehicle *before* becoming alert, and then escapes considerably *after* becoming alert can help choose the right model. If a species consistently detects the approaching vehicle much earlier than escaping, the perceptual limits hypothesis would have limited application generating quantitative predictions because it violates one of its key assumptions (Fig 6). Along similar lines, if a species consistently detects the approaching vehicle much earlier than becoming alert, the FEAR hypothesis and the Bayesian optimal escape model would be limited in their ability to generate quantitative predictions because of violations of the AD = DD assumption (Fig 6). It is likely that the timing of detection, alert, and escape responses is a function of the sensory constraints and metabolic needs of different species; which could lead to species-specific differences. Unfortunately, the empirical evidence to elucidate these questions is scant.

### Implications for reducing animal-vehicle collisions

Escape rules (temporal margin of safety, spatial margin of safety, delayed margin of safety) appear to have an important role in affecting the predictive ability of models of escape behavior. Yet, there are few studies that characterized the escape rules different species use relative to high-speed approaching vehicles. Empirical studies manipulating approach speed with real or virtual vehicles found that brown-headed cowbirds [15], turkey vultures [14], and rock pigeons [16] appear to follow the spatial margin of safety, at least at lower speeds, with inconclusive results for white-tailed deer [99] .

Species that follow a spatial margin of safety are particularly vulnerable to vehicle collisions because they do not adjust their avoidance responses to the unnaturally high speeds of modern vehicles. As a result, they often do not allow sufficient time to clear the path of the oncoming vehicle once they initiate their avoidance response. Furthermore, human safety research using virtual vehicles provides empirical support for the higher risk incurred when following a spatial margin of safety to avoid vehicles [100–102]. Humans whose crossing decisions are based on distance from the approaching vehicle (i.e., a spatial margin of safety) are more likely to be struck by the virtual vehicle when crossing a street than those whose street crossing decisions are based on the estimated time until the vehicle impact (i.e., a temporal margin of safety) [100, 103].

For species following the spatial margin of safety, we propose to calculate the vehicle speed at which an animal can no longer avoid a collision with an oncoming vehicle, even when the vehicle is detected and continuously monitored. We call this metric the Critical Vehicle Approach Speed (CVAS). To calculate CVAS, estimates are needed for the threshold FID used to escape consistently irrespective of vehicle speed (as described above) as well as the time needed for escape (i.e., to clear the path of the oncoming vehicle and avoid a collision) after the avoidance response is initiated. As an example, we estimate CVAS (expressed in km/h) for the brown-headed cowbird based on data provided by DeVault et al. (2015) [15]. In that study, cowbirds maintained a flight initiation distance (FID) of 28 m for vehicles approaching at speeds of ≤150 km/h, and required 0.80 s (t) to avoid a collision once an avoidance maneuver was initiated (3 m; roughly the width of one lane in a standard road). We calculate CVAS = FID / (t × 0.2778) = 126 km/h. We conclude that brown-headed cowbirds have a high risk of colliding with a vehicle that is traveling at a constant speed ≥126 km/h, even under ideal conditions for that species to detect and track the vehicle (S4).

The CVAS concept has implications for management of populations that are especially susceptible to road mortality. Again, we emphasize that this metric requires data on threshold FID relative to vehicle approach and escape time per species of interest. CVAS could be used as a guide to establish vehicle speed limits in areas where species of conservation concern are located (assuming those species follow a spatial margin of safety). To illustrate, consider a fictional bird species that commonly loafs at the end of an airport runway and does not respond well to dispersal via non-lethal hazing. Prior research determined that the species follows a spatial margin of safety with a consistent FID of 70 m across a range of vehicle speeds, and requires 2.1 s to successfully clear the path of an approaching aircraft once the avoidance behavior is initiated. CVAS is calculated at 120 km/h. However, aircraft on that runway typically reach 240 km/h at the point of takeoff, creating conditions that result in a high likelihood of a bird-aircraft collision. Researchers developing mitigation methods such as onboard aircraft lights designed to enhance perceived risk and stimulate earlier avoidance behaviors [105–107] could target an increase in the flight initiation distance for that species from 70 m to at least 150 m as a goal with aircraft lighting development and experimentation. Consequently, the CVAS provides a target effect size to increase FID to reach a CVAS = 257 km/h, greater than the speed of the oncoming aircraft, which would greatly reduce the likelihood of collision. A similar rationale can be applied to interstate highways, train tracks, and other travel corridors for vehicles where it might be impractical to reduce typical vehicle speeds and where other mitigation methods designed to enhance avoidance behaviors can be applied (i.e., lights, sounds, etc.).

There are, however, some limitations in the application of the CVAS concept. First, the escape rules of too few species have been evaluated empirically, as summarized above. Second, escape time (the amount of time needed to successfully avoid the collision) is also poorly studied, although it could be estimated based on movement speed, body mass, or other physical measurements. Third, a spatial margin of safety might be followed by a species only up to some threshold speed, above which a different avoidance strategy (or none at all) might be followed [15] . Future research is needed to determine how widespread the seemingly maladaptive use of a spatial margin of safety for vehicle avoidance is across species that are commonly struck by vehicles, and how and why risk assessment strategies vary across species. Understanding such differences would allow researchers to identify trends in risk assessment and avoidance strategies related to certain taxa, life histories, and environmental variables.

### New models of escape behavior to high-speed vehicles are necessary

One of the most important theoretical implications of our findings is that new models of escape behavior tailored to high-speed vehicles (instead of predators) are necessary to improve our ability to make quantitative predictions. However, there is a lot of empirical groundwork required to start developing such new models because of the large gap in our understanding of how animals detect, become alert, and eventually escape from high-speed vehicles.

An immediate challenge is to understand the mechanisms underlying cue detection and processing, functions that inform antipredator behavior [108]. We contend that mechanisms informing escape decision-making must consider the sensory perspective [4,15,74]. Some research has focused largely on the visual sensory path relative to terrestrial bird and mammal responses to vehicle approach [109], and auditory cues relative to marine mammal responses to boat traffic [110–111]. The effectiveness of incorporating a sensory ecology approach to the new models of escape behavior will require understanding of at least two components: configuration of the sensory path relative to high-speed object detection and response, how cues are processed once detected at far distances, and the behavioral repertoire used in escape decisions (whether adaptive or maladaptive). Additionally, different species may vary their response due to different life-history traits. Consequently, it is essential to determine the degree to which the sensory, cognitive and anatomical capacity of different species may account for differential responses to high-speed vehicles. It is highly possible that different species have threshold speeds above which their capacity to avoid collisions is basically overwhelmed [4]. Clearly, there is evidence that animals can adapt to vehicle-associated risk [112, 113] but we will not understand how adaptations succeed or fail, or how to mitigate against failure, absent first identifying the mechanism(s) governing responses.

Future models should also consider individual differences in behavior and learning. Individual differences in escape responses maybe attributed to an animal’s state, previous experience, or the configuration of their sensory systems [11,17,114–118] . Future models should account for behavioral variation in escape distance within an individual (i.e., intraindividual) and between individuals (i.e., interindividual). Behavioral responses to perceived risk (e.g., such as vehicle approach), if governed largely by intraindividual differences, are repeatable [119–123] .

However, if individual responses vary to the same types of stimuli such that intraindividual variation is greater than interindividual variation, then escape behavior will not be repeatable [11]. Intraindividual and interindividual differences may result in differences in survivability when approached repeatedly by a high-speed vehicle. Exploring the effect of individual differences on escape behavior may help improve the resolution of future models but also may provide key insights about how individual differences could potentially be exploited to mitigate vehicle collisions.

Implicit to the role of learned escape behavior is that an animal, through experience, realizes some benefit in its response to a particular stimulus [19,26,124]. For example, watching a conspecific that is stalked, attacked, and killed by a predator can contribute to future perceived predation risk [20, 125] . Further, behavioral plasticity in response to predation risk, by which an animal might “manage” undue interruption in foraging, etc., can be affected by prior experience with predators [126–129] . We have very little understanding of these dynamics relative to high-speed vehicle risk.

Finally, future models should seriously consider incorporating the effects of learning, as it can play a key role in conservation and management. Habituation is defined as a decrease in behavioral response to repeated stimulation, not due to sensory adaptation or fatigue [130]. In contrast, sensitization is an enhancement of behavioral response to repeated stimulation [131]. The context of exposure (e.g., resources available, group size, presence of young), frequency of exposure [17, 131], as well as negative experiences that might influence learning, can influence how long habituation or sensitization might be shown in response to particular cues [131]. In addition, social information might override personal information when it comes to escaping from an approaching vehicle [10]. Incorporating learning in future models has the potential to greatly increase their applicability to real management scenarios to reduce animal-vehicle collision risk. Nevertheless, we acknowledge that developing future models will take extensive effort and time (and funding availability), but we believe it is a fruitful area of research that can converge the interests of theoretical and applied ecologists.

## Acknowledgements

We would like to thank Jeff Lucas, Ximena Bernal, Catherine Searle, and Benny Goller for feedback on the review. This review was in part funded by the United States Department of Agriculture, Animal and Planth Health and Inpsection Service, and the Federal Aviation Administration. Contributions by T.L. DeVault were partially supported by the U.S. Department of Energy under award #DE-EM0004391 to the University of Georgia Research Foundation.

## Notes

https://osf.io/b4cs2/?view_only=94c33be8a061462399c81dace7884151

https://github.com/ryanlunn/AntipredatorTheoryAndVehicleSpeed

### Competing Interest Statement

The authors have declared no competing interest.

